# A model of rapid homeostatic plasticity accounts for hidden, long-lasting changes in a neuronal circuit after exposure to high potassium

**DOI:** 10.1101/2021.07.01.450770

**Authors:** Mara C.P. Rue, Leandro Alonso, Eve Marder

**Affiliations:** Biology Department and Volen Center, Brandeis University, Waltham, MA 02454

**Keywords:** *Cancer borealis*, neuronal oscillators, stomatogastric ganglion, homeostasis

## Abstract

Neural circuits must both function reliably and flexibly adapt to changes in their environment. We studied how both biological neurons and computational models respond to high potassium concentrations. Pyloric neurons of the crab stomatogastric ganglion (STG) initially become quiescent, then recover spiking activity in high potassium saline. The neurons retain this adaptation and recover more rapidly in subsequent high potassium applications, even after hours in control saline. We constructed a novel activity-dependent computational model that qualitatively captures these results. In this model, regulation of conductances is gated on and off depending on how far the neuron is from its target activity. This allows the model neuron to retain a trace of past perturbations even after it returns to its target activity in control conditions. Thus, perturbation, followed by recovery of normal activity, can hide cryptic changes in neuronal properties that are only revealed by subsequent perturbations.

## Introduction

An enigmatic property that all nervous systems share is their ability to maintain proper physiological function despite ongoing perturbations to their activity and constant turnover of their ion channels and other cellular components. At the same time, neural circuits must be able to adapt to varying internal and external environments.

For all organisms, maintaining the appropriate ionic composition of the extracellular milieu is critical for normal physiological function, and the potassium gradient is particularly important for the maintenance of resting membrane potential and normal activity levels. It is therefore unsurprising that altered potassium homeostasis occurs in a wide array of conditions including heart disease, kidney failure, thermal stress, epilepsy, traumatic brain injury and stroke ^1–7^. In addition to these pathological disease states, altered extracellular potassium levels are routinely used by researchers as a physiologically relevant depolarizing stimulus to increase neuronal activity or as a proxy for excitatory inputs ^8–11^. Nonetheless, many studies employing high potassium do not record the physiological response of neurons. Those that do record physiologically often look only at long-term, chronic changes of populations of neurons over days to weeks^12–14^. But changing extracellular potassium concentration will immediately affect neuronal membrane potentials, and thus may activate rapid adaptation mechanisms. Given this, we were interested to observe how elevations in extracellular potassium levels would affect individual neurons over time.

By studying how extracellular potassium concentrations affect identified neurons, we have an opportunity to observe mechanisms of adaptation to a global depolarization. Typically, researchers classify activity-dependent adaptation into several distinct timescales. The shortest activity-dependent adaptation processes such as spike frequency adaptation emerge from ion channel properties that occur on the millisecond timescale. Over longer timeframes, activity-dependent homeostatic mechanisms actively regulate ion channel expression and synaptic weights to maintain stable function in the face of physiological perturbation^15–25^. These homeostatic processes are commonly thought to act over hours to days and require protein synthesis. However, similar feedback mechanisms can also drive more rapid adaptation over intermediate timescales on the order of minutes. For instance, changes in *effective* conductance density can occur quickly through phosphorylation of ion channels^26–28^ or rapid insertion of ion channels^29^.

Models of activity-dependent plasticity or homeostasis generally involve feedback mechanisms that monitor internal calcium dynamics to modify the conductance densities of specific ion channels. Using these rules, one can build neurons with given target activities that can recover from perturbation ^23, 30^. However, all current models of homeostatic plasticity have some limitations. For instance, in models using a single calcium sensor, neurons can be robust to some perturbations, but vulnerable to targeted deletion or changes in specific conductances^31^. Conversely, models involving more than one calcium sensor can be inherently unstable^23, 30^. Finally, conventional computational models of neurons are far more vulnerable to perturbation than biological neurons ^20, 23, 30, 32–34^. This suggests that some mechanisms of activity-dependent adaptation must be included in computational models to study how neurons respond to perturbations.

The crustacean stomatogastric ganglion (STG) is an excellent model system in which to study underlying network dynamics and mechanisms of circuit robustness both through recording from well-studied identified neurons and computational models of those neurons^33, 35–38^. Importantly, the physiological behavior of each neuron within the STG is relatively stereotyped, allowing us to determine whether a given pattern of activity is normal. This system therefore provides an excellent paradigm in which to study how a neural circuit can achieve stable adaptation to global perturbation while maintaining its characteristic physiological function. Taking advantage of this tractable and well-defined system, we investigated the response of neurons to high potassium and describe a case of intermediate-term (minutes) adaptation to a global perturbation which is retained over long time periods (hours). We then used these observations to modify a computational model of homeostatic adaptation. These studies demonstrate a mechanism by which adaptation can lead to cryptic changes in neuronal excitability that become visible only in response to a subsequent environmental challenge.

## Results

### Short-term adaption of pyloric neurons to elevated potassium concentrations

The pyloric central pattern generator within the STG drives filtering of food particles through the foregut *in vivo* ^39^. The same network activity persists *in vitro* ^40^ and can be monitored using a combination of intracellular and extracellular recordings. The pyloric network is driven by the anterior burster (AB) neuron together with the two pyloric dilator (PD) neurons, which together form a pacemaker kernel. In this study we focused on the regular bursting activity of the PD neuron as a proxy for robustness of the pyloric circuit (Fig. 1a(*i*)). For all experiments, the stomatogastric nervous system (STNS) was dissected intact from the stomach of the crab, *Cancer borealis,* and pinned in a dish, allowing us to change the composition of continuously superfused saline.

**Figure 1:**
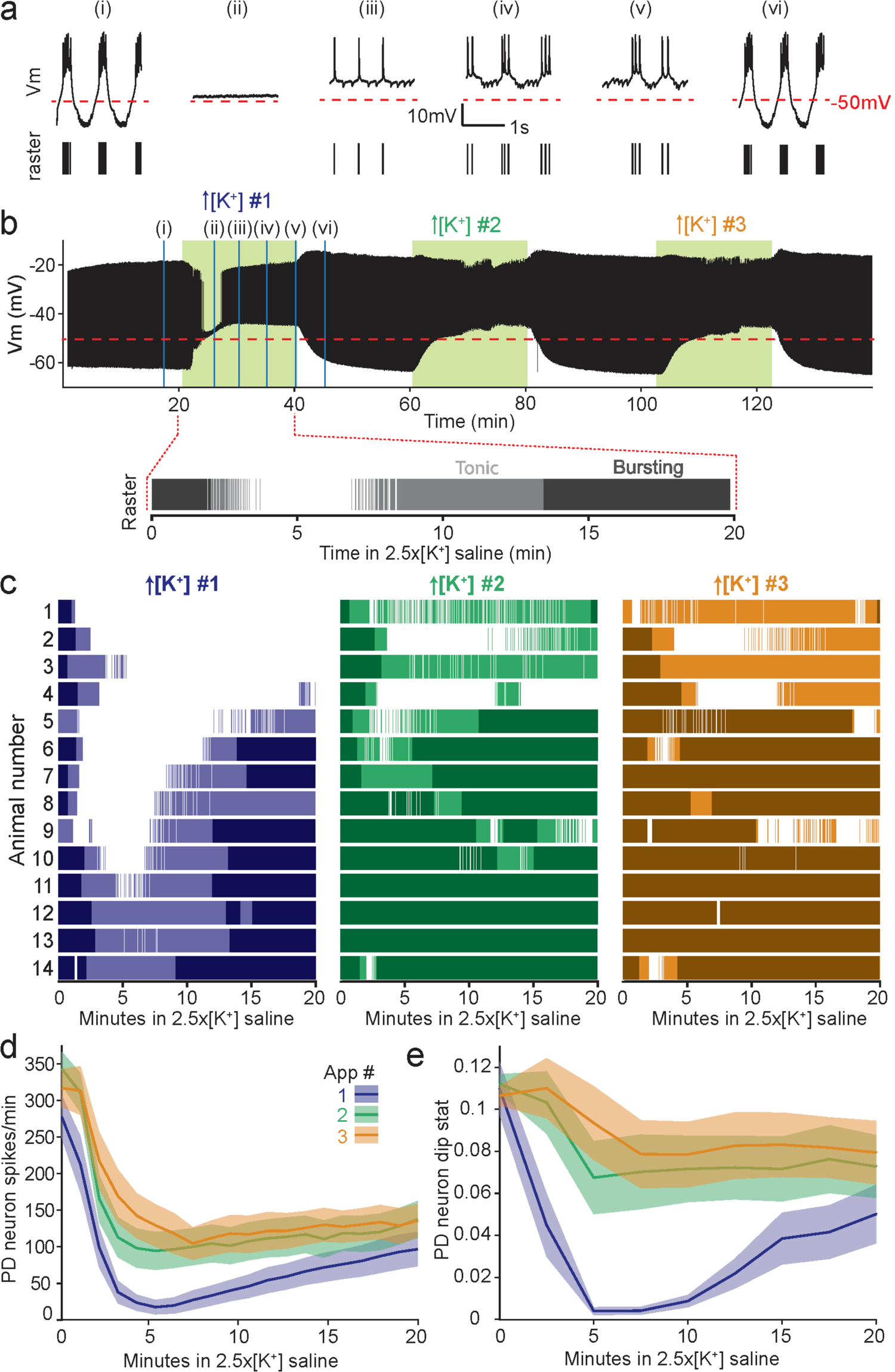
PD neurons adapt to elevated potassium concentrations and are more robust to the perturbation upon repeated exposure. **(a)** Two-second segments of a PD neuron’s activity in control physiological saline (*i*), five (*ii*), ten (*iii*), fifteen (*iv*) and twenty (*v*) minutes into the first application of 2.5x[K^+^] saline and during the first wash period (*vi*). Below each trace is shown the spike raster with a vertical line plotted for every action potential in the trace. **(b)** Voltage trace for the same PD neuron over the entire experiment. Green shaded boxes indicate time of 2.5x[K^+^] saline superperfusion. Below this trace is shown is a raster plot of spiking activity for the entire first application of 2.5x[K^+^] saline, with bursting activity plotted in a darker shade and tonic firing plotted in a lighter shade. **(c)** Raster plots of spiking activity in 2.5x[K^+^] saline for fourteen PD neurons exposed to three repeated exposures. For all plots, bursting activity is plotted in a darker shade and tonic firing in a lighter shade. **(d)** Average PD spikes per minute for all three applications are plotted in the dark line with SEM shaded regions around them. **(e)** Average PD dip value for all three applications are plotted in the dark line with SEM shaded regions around them.

We previously demonstrated that pyloric neurons depolarize, temporarily become silent in high potassium saline, and subsequently recover spiking activity through a change in cell-intrinsic excitability ^38^. In this work we studied repeated applications of high potassium to ask if neurons retain a long-term trace or memory of this adaptation. When PD neurons are first exposed to 2.5 times the physiological concentration of extracellular potassium (2.5x[K^+^] saline), the neuron depolarizes and becomes quiescent (Fig. 1a*ii*) before recovering spiking and later bursting activity over 20 minutes in elevated extracellular potassium (Fig. 1a*iii*-*v*). This change in activity can be visualized by the raw voltage traces (Fig. 1a, top) and simple raster plots where a line is plotted for each action potential in the respective PD neuron (Fig. 1a, bottom).

We superfused the STNS with three 20-minute 2.5x[K^+^] saline exposures interspersed with 20-minute washes in physiological (control) saline (Fig. 1b). Repeated exposure to elevated extracellular potassium resulted in shorter or diminished periods of quiescence and more robust PD neuron spiking activity compared to the initial application (Fig. 1b, c). In all animals (N=14), PD neurons exhibited more spiking and bursting behavior in high [K^+^] applications #2 and #3 compared to the first application (Fig. 1d, Friedman’s test, Q(2) = 23.57, multiple comparisons with Bonferroni correction. The number of spikes during the first application differs from second and third for minutes 4 – 13 after beginning of application (p <0.0025 for all)). Nonetheless, significant individual variability can also be observed across animals.

Under normal physiological conditions, pyloric neurons produce bursts of action potentials, which are necessary to drive rhythmic contractions of muscles within the stomach of the crab^41^. Therefore, we also characterized the “burstiness” of pyloric neurons during exposure to high potassium saline using Hartigan’s dip statistic, in which higher numbers indicate more burst-like activity. For all PD neurons, the dip statistic was higher throughout the second and third high potassium applications compared to the first (Fig. 1e, Friedman’s test, Q(2) = 16.87, multiple comparisons with Bonferroni correction. Dip value during the first application differs from second and third for minutes 6 – 12 after beginning of application (p <0.005 for all)). Overall, the improved spiking activity and “burstiness” of PD neurons in high potassium saline upon repeated applications indicates that the intrinsic properties of pyloric neurons are altered by a single exposure to high potassium, and that these changes are maintained after 20-minute washes in control saline.

### Pyloric activity in control saline is unchanged following potassium perturbation

Given that pyloric neurons rapidly adapt to the high potassium perturbation, we might expect that this change in excitability would affect the neurons’ overall activity level. To see if this was the case, we directly compared the bursting activity of each PD neuron in control saline and after each high potassium application (Wash #1-3). Figure 2a depicts example traces from PD neurons from three preparations. Here the animal number to the left corresponds to the animal numbers shown in Figure 1. Although all the PD neurons shown here had distinct sensitivities to high potassium saline (see Fig. 1c), the baseline activity of the neurons is similar across animals. Additionally, within each preparation the activity in the washes appears similar to baseline. For all PD neurons, we analyzed the bursting activity in the last ten minutes of baseline and washes #1-3. The burst frequency of PD neurons was unchanged in control saline regardless of the wash number (Fig. 2b, Friedman’s test, Q(3) = 2.45, p = 0.46). Similarly, there was no change in the average number of spikes per burst (Fig. 2c, Friedman’s test Q(3) = 4.66, p = 0.17). In summary, although PD neurons show robust adaptation to high potassium saline, we observe no differences in bursting behavior under control conditions.

**Figure 2:**
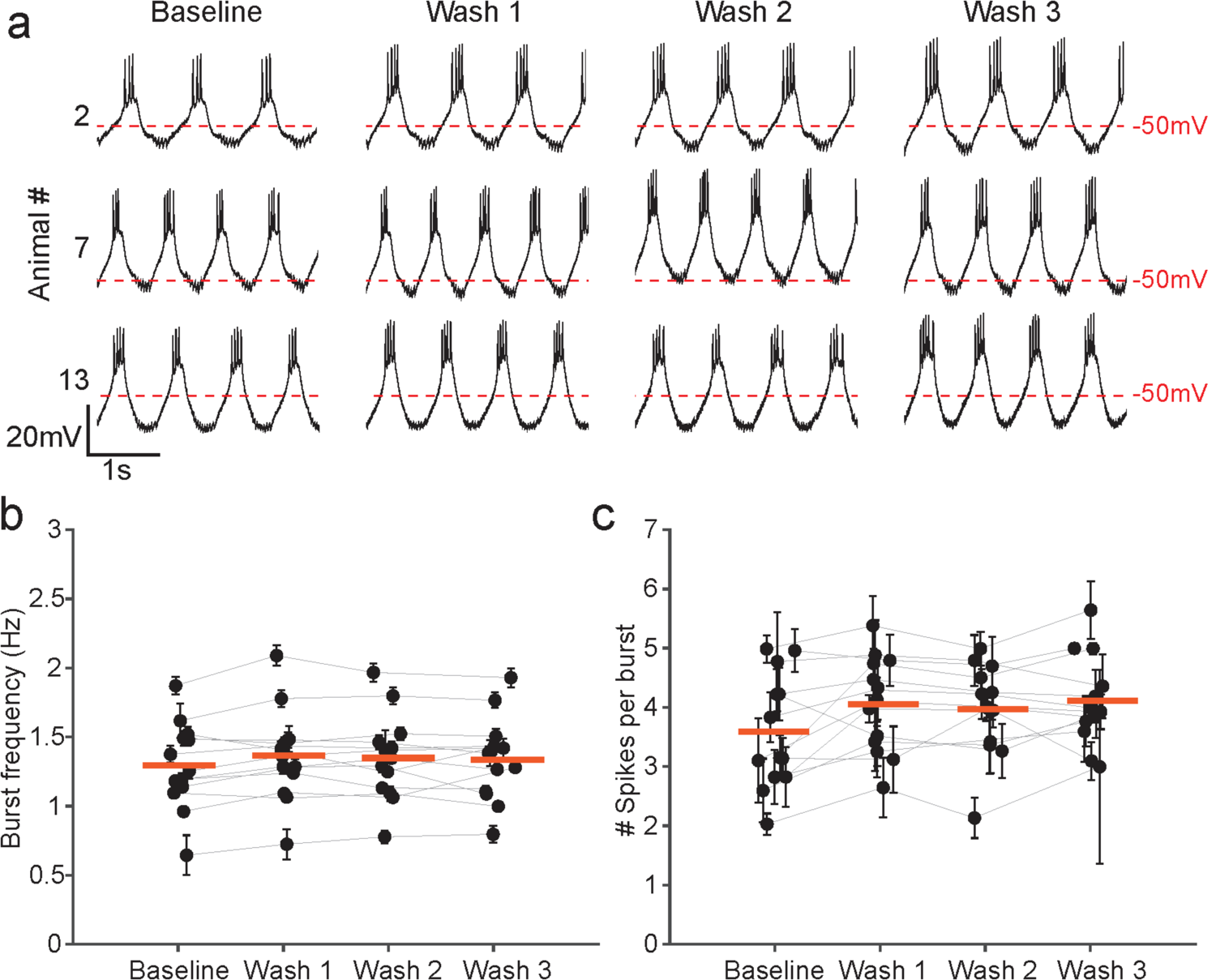
Bursting activity of PD neurons in control saline is unchanged after high potassium applications. **(a)** Three-second segments of three PD neurons’ activity in baseline, wash #1, wash #2, and wash #3 after high potassium applications. All traces are in control saline with normal physiological potassium concentration. The animal numbers on the left correspond to the animal numbers in Figure 1C. **(b)** Average burst frequency in each condition for all PD neurons with error bars representing standard deviation. Individual experiments are connected with light grey lines. The mean of all PD burst frequencies for each time point is indicated by a thick red line. **(c)** Average spikes per burst in each condition for all PD neurons; error bars represent standard deviation. Individual experiments are connected with light grey lines. The mean of all PD spikes per burst for each time point is indicated by a thick red line.

### Adaptation to elevated potassium is maintained long-term after several hours in control saline

Because our potassium applications are relatively brief, one might expect a PD neuron to return to their baseline sensitivity after a period of time under control conditions and lose the enhanced robustness to high potassium saline.

To test this, we performed additional experiments in which we applied the same three rapid 20-minute applications of 2.5x[K^+^] saline interspersed with 20-minute washes in control saline, followed by a three-hour wash, and finally a fourth 20-minute 2.5x[K^+^] saline application. Here, unlike in the previous set of experiments, the third wash is many times longer than the perturbation that drove the change in robustness. Again, PD neurons showed improved robustness over the first three applications of high potassium saline (example traces of activity at 15 minutes in 2.5x[K^+^] saline, Fig. 3A*ii, iii, iv*). After the three-hour wash in control saline, the representative PD neuron (animal 15) shown in Figure 3a maintained and improved this robust response to 2.5x[K^+^] saline (Fig. 3a*v*, 3b). All PD neurons in this set of six experiments retained their decreased sensitivity to high potassium saline after extended wash (Fig. 3c). Overall, the number of spikes per minute in 2.5x[K^+^] saline increased across the first three applications, and was maintained in the fourth application after the extended wash (Fig. 3d, Friedman’s test Q(3) = 18.54, multiple comparisons with Bonferroni correction. The number of spikes per minute during the first application differs from second, third and fourth for minutes 2 – 14 after beginning of application (p <0.0025 for all)). PD neurons exhibited more bursting activity in high potassium saline in applications #2-4 compared to the first (Fig. 3e, Friedman’s test Q(3) = 10.39, multiple comparisons with Bonferroni correction. Dip statistic during the first application differs from second, third and fourth for minutes 2 – 12 after beginning of application (p <0.005 for all)). Thus, pyloric neurons retain an imprint of past exposures to high potassium saline, even after a wash period much longer than the perturbation itself and despite the fact that unperturbed recordings show little overt sign of this adaptation.

**Figure 3:**
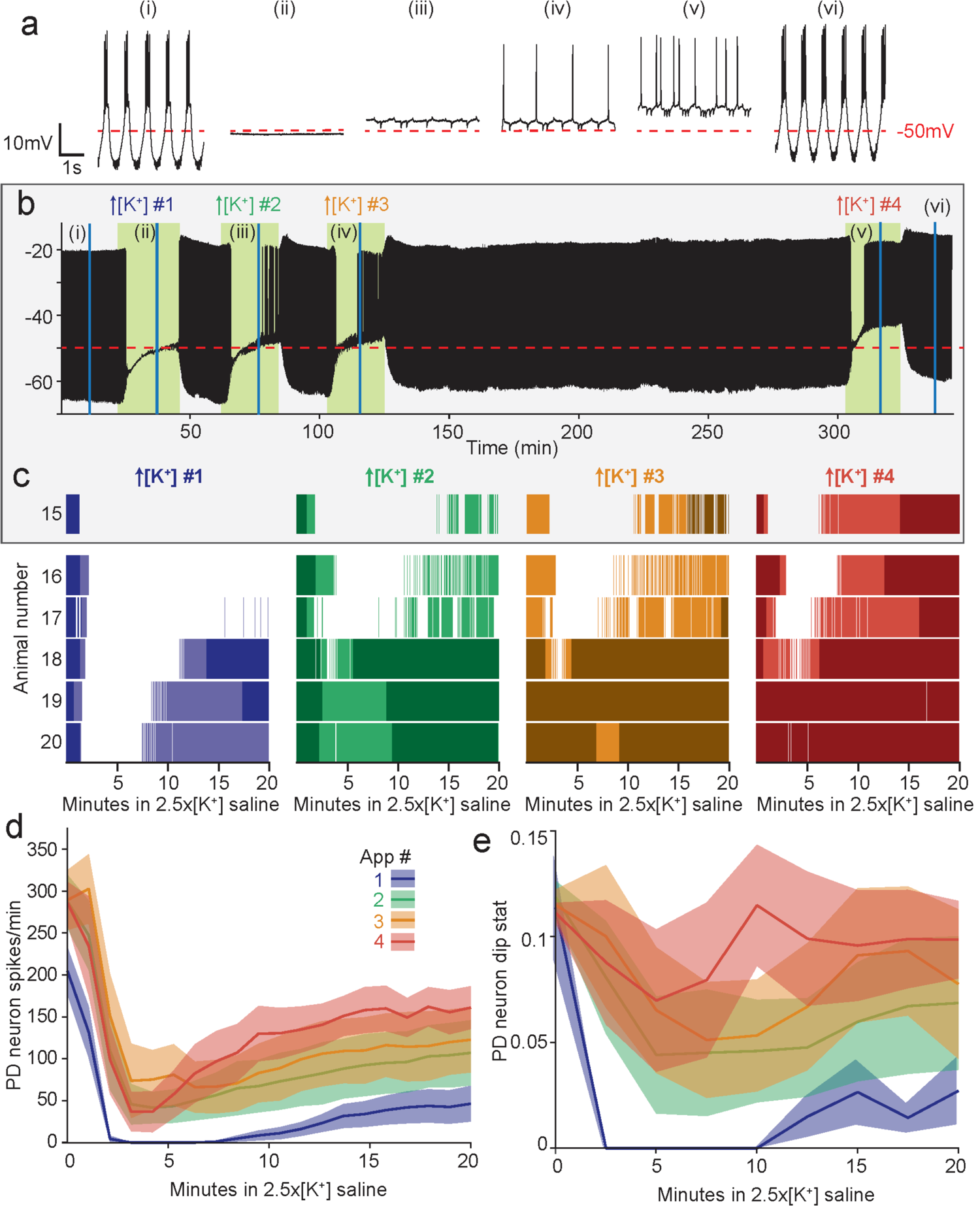
PD neurons retain adaptation to high potassium saline even after several hours of wash in control saline. **(a)** Four-second segments of a PD neuron’s activity in control physiological saline (*i*), and at fifteen minutes into the first (*ii*), second (*iii*), third (*iv*), and fourth (*v*) applications of 2.5x[K^+^] saline, and upon the final wash in control saline (*vi*). **(b)** Voltage trace for the same PD neuron over the entire experiment. Green shaded boxes indicate time of 2.5x[K^+^] saline superperfusion. Below this trace is shown is a raster plot of spiking activity for each of the four applications of 2.5x[K^+^] saline, with bursting activity plotted in a darker shade and tonic firing plotted in a lighter shade. **(c)** Raster plots of spiking activity in 2.5x[K^+^] saline for six PD neurons (15-20) exposed to the same four repeated exposures. For all plots, bursting activity is plotted in a darker shade and tonic firing in a lighter shade. The top raster (15) is the same animal as that shown in **a** and **b** above **(d)** Average PD spikes per minute for all four applications are plotted in the dark line with SEM shaded regions around them **(e)** Average PD dip value for all three applications are plotted in the dark line with SEM shaded regions around the lines.

### Modeling bursting neurons exposed to high potassium

We constructed a computational model of a neuron that captures the main qualitative observations in the previous experiments, and which reveals features of adaptation mechanisms that are difficult to see directly. To this end, we evaluate how several models with different long-term regulation properties respond to the same high potassium perturbation. These include (a) a conventional conductance-based model with no regulation, (b) a three-sensor homeostatic model modified from Liu et al (1998)^30^, and (c) a new three-sensor homeostatic model with novel conductance regulation properties. Figure 4 shows simulations in which we applied the high potassium perturbation (shifted E_K_ from −80mV to −40mV) to the three different model neurons in intervals of 20 minutes. We also simulated a long wash period of 3 hours, followed by a final 20-minute application of high potassium, similar to the experiments in Figure 3. For all panels, the membrane potential is shown on top, and the conductance densities of the currents are shown below. The numerals below the voltage trace indicate the type of activity pattern (Fig. 4*i*-*iv*) at different temporal segments.

**Figure 4:**
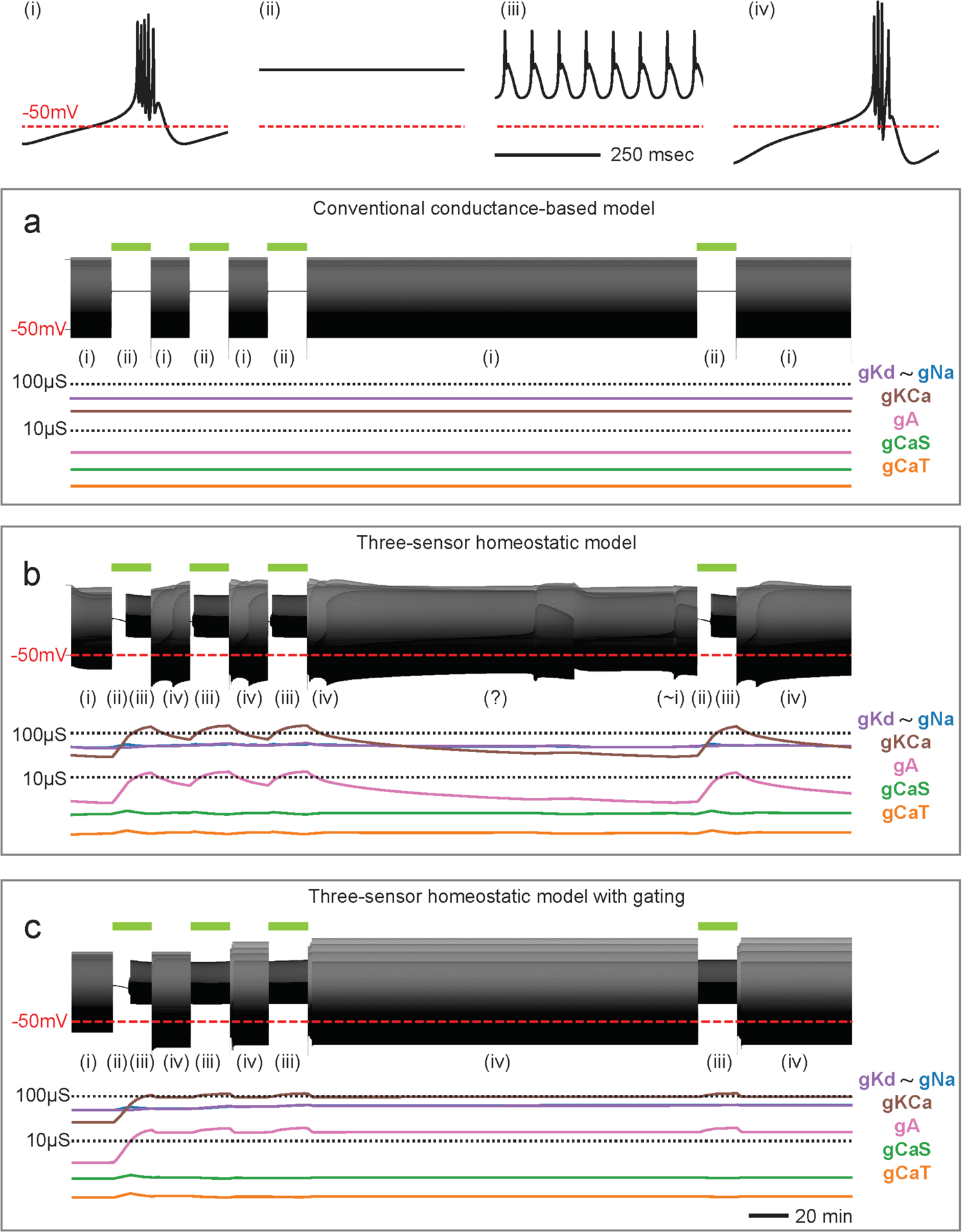
Model bursting neuron response to high potassium. The top panels show representative voltage traces (*i*-*iv*) for all models **(a-c)** and green bars above the voltage trace represent the high potassium perturbation. The compressed voltage trace of the model neuron is shown in the top panel and the evolution of that model’s conductance densities are shown below **(a)** The model does not have a regulation mechanism, and the conductances are fixed. The model becomes quiescent in the high potassium condition regardless of its history. **(b)** The model regulates its conductances in an activity dependent manner to stabilize the control bursting pattern. The model becomes quiescent in high potassium but recovers spiking over ten minutes. During the long wash, conductances return to the control values, and history dependance is erased. **(c)** The model is identical to **b**, with an additional feedback signal (Sf) that monitors if the cell is bursting or not. The model regulates its conductances only if the feedback signal is low. The conductances stay constant during the long wash because the cell is bursting, and the feedback signal turns off the regulation mechanism.

### The model neuron becomes quiescent in the first exposure to high potassium

Figure 4a depicts a conventional conductance-based neuronal model. In this model, when the potassium reversal potential is changed the membrane potential depolarizes and the cell becomes quiescent. The model remains quiescent during the high potassium condition, recovers bursting activity in wash, and unsurprisingly, the model does not adapt and becomes quiescent again when subsequently exposed. This simulation strongly suggests that to replicate the experimental data the conductance densities in the model neuron must change. Note that the maximal conductances in this model do not change over time.

### A homeostatic model with bounded current densities can rapidly adapt to the high potassium perturbation, but does not retain a long-term memory

This led us to revisit a family of models with homeostatic regulation that have been used for many years to understand how neurons develop proper bursting behavior and adapt to changes in the environment ^20, 30, 34, 42^. We devised a modification of the model by Liu et al. ^30^ which uses three sensors (fast, slow and DC filter of the calcium current) that monitor calcium currents and employs them to modify the neuron’s effective conductance densities and achieve a target activity. The model can be expressed as follows,

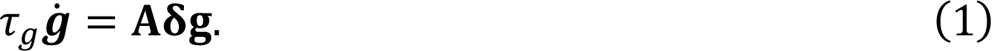

Here **A** is a fixed matrix that defines how the sensor outputs translate into conductance changes. δ(t) is a vector that measures how close each sensor is to its set point (see methods). Ideally, the conductances stay constant when all components δ_i_ = 0, which occurs when each sensor is at its set point. This model can recover from several perturbations including changes in the reversal potential of potassium currents ^43^.

However, one drawback of model as implemented in Liu et al. ^30^ is that if all three sensors are not satisfied at the same time, this can result in run-away activity which leads conductances to increase rapidly, causing it to diverge ^30, 43^. In other words, over long time periods model (1) will often become unstable. Despite this limitation, the three sensors are useful in distinguishing between different patterns of activity. For example, in the case of a neuron with a periodically bursting target activity, a perturbation could switch the activity to a tonic spiking state. For the cell to recover back to the bursting state, it must be able to sense a difference between the bursting state and the tonic spiking state. As Liu et al. ^30^ shows, when using only one calcium sensor it is not always possible to tease apart these two activity patterns because the average calcium in the model cell can be similar in both regimes. In this situation the conductances would stay constant and the model neuron would not recover from such a perturbation. The high extracellular potassium perturbation studied here runs into the same difficulty: the average calcium levels in the cell during bursting in control and the quiescent state in high potassium saline are similar. Therefore, we sought a modification of the model that would allow it to operate in a stable fashion with the multiple sensors needed to distinguish tonic spiking from bursting activity.

Hence, we incorporated explicitly in this model the biological assumption that conductances can’t grow indefinitely and must be bounded by some maximum value. This modification prevents the model from diverging, but preserves many of its properties such as the possibility of recovering spiking during the high potassium condition,

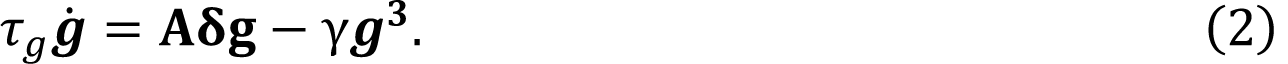

With this modification, model (2) will respond to perturbations in a similar way as the original formulation (γ = 0) unless some conductances are too large. If **g** is too large, the cubic term will dominate and ġ < **0**, meaning that **g** will decrease. The parameter γ can be used to set a bound for how much a conductance can grow. We chose a cubic term for simplicity but note that any function that satisfies ġ < **0** for **g** sufficiently large would prevent the model from diverging.

Model (2), Figure 4b, retains a transient memory of prior adaptation to high potassium saline. There are multiple regions in conductance space that correspond to bursting patterns under control and wash conditions, but our recovery mechanism favors some regions over others because of the details of the control scheme and other parameters (See Methods) ^43^. When model (2) is returned to the control condition after the first high potassium exposure (wash), the values of the conductances and the bursting waveform are slightly different from those before the first exposure (compare Fig. 4*i* and 4*iv)* As time unfolds in control saline, the conductances trend back to their starting values. Because of our chosen timescale for conductance changes, τ_%_ = 2 minutes, over the 20-minute wash period the conductances do not reach their original set-points. Therefore, over the first three applications of high potassium saline in Figure 4b, the neuron is more robust to the second and third application compared to the first. Nevertheless, the conductances will trend back to their starting values if given sufficient time, as happens in the long three-hour wash period. For this reason, in Figure 4b the response to high potassium in the fourth application of high potassium is akin to the first response. This contrasts with the biological data showing that PD neurons can maintain robustness to the high potassium perturbation over a long wash period (Fig. 3).

### Addition of a novel activity-dependent gating mechanism for homeostatic plasticity allows the model neuron to retain long-term memory of past perturbation

To allow the model neuron to retain its adaptation to previous perturbations, we next wanted to enforce the condition that the bursting patterns in control (Fig. 4*i*) and in wash (Fig. 4*iv*) are equally acceptable, and that the cells’ conductances need not drift back to their starting values. In the model by Liu et al. ^30^, the readings from the three sensors are used to drive changes in conductances; equilibrium is expected when the sensors are simultaneously at their set points. In a sense, the sensors in this model are playing a dual role: they drive changes in conductances in a specific way, and they also monitor that the cell is at its target pattern, because the equilibrium condition (ġ = **0**) requires all sensors to be at their set points. Here we explored a new modification: that the specific way in which the model modifies its conductances is independent of the equilibrium condition. In this way, we incorporate the possibility that there are two pathways: one that drives changes in the conductances (possibly but not necessarily using sensor readings), and another that controls whether the regulation mechanism is active or not. For this we hypothesized that there is a feedback signal that combines the readings of the sensors, and that this signal modulates the timescale of conductance regulation. We implemented this idea using a state variable α that takes values between zero and one. If the model is bursting periodically, the feedback signal is high and α → 0. If the feedback signal is low and the model’s activity is other than the target pattern, then α → 1.

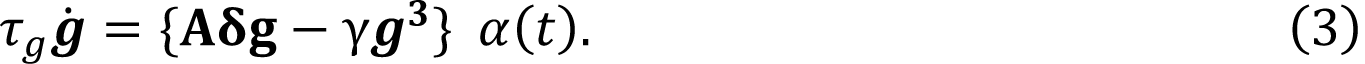

Figure 4c shows a simulation of model (3) subjected to the same experimental paradigm as before. In control conditions the feedback signal is high (α ≈ 0), so the conductances stay constant until the first exposure to high potassium. When the cell becomes quiescent, the feedback signal is low (α ≈ 1), so the recovery mechanism is activated similarly to Fig. 4b and allows the cell to recover spiking activity in high potassium saline. As before, the cell recovers bursting upon wash, but now the feedback signal is high, and the recovery mechanism is turned off (α ≈ 0). Instead of returning to their control values the conductances stay constant during the wash intervals regardless of the duration of the wash period. In this way, model (3) will remain robust to the high potassium saline application after a long wash period, and for this reason response to perturbation in application four is different from that in the first application.

### Time course of recovery in high potassium depends on starting conductance densities

In our experiments, the amount of time it takes for PD neurons to recover spiking activity upon the first exposure to high potassium saline varies widely across animals; some neurons regain spiking almost immediately while others remain silent for almost 20 minutes (Fig. 1c). In identified neurons from the STG, mRNA copy number for ion channels and recorded currents can vary 2 to 6-fold accross individuals ^44–48^. Therefore, one way to account for the variability in recovery time is to assume that individual differences between PD neurons determine the sensitivity to high potassium saline. To test this hypothesis, we used our newly devised model (3) to investigate whether individual differences in conductance density between neurons may be sufficient to explain the observed variability.

We generated 9 model neurons that use the same adaptation mechanisms as model (3). Previous studies have demonstrated that model neurons with different underlying parameters can nonetheless have similar activity patterns^23, 33, 49^, and we replicate these findings here. Figure 5 shows the response of five representative model neurons (models P, Q, R, S and T) to the high potassium perturbation. The example traces show the membrane potential of the models in control conditions (Fig. 5a*i*) and at ten minutes into the first high potassium exposure (Fig. 5a*ii*). Note that all models exhibit similar bursting patterns of activity, although each has a different set of starting conductance densities (Fig. 5a*i*, control conditions). The compressed membrane potential traces for each model are shown in Figure 5b. All five models become quiescent immediately after exposure to high potassium saline recover spiking activity after a variable amount of time.

**Figure 5:**
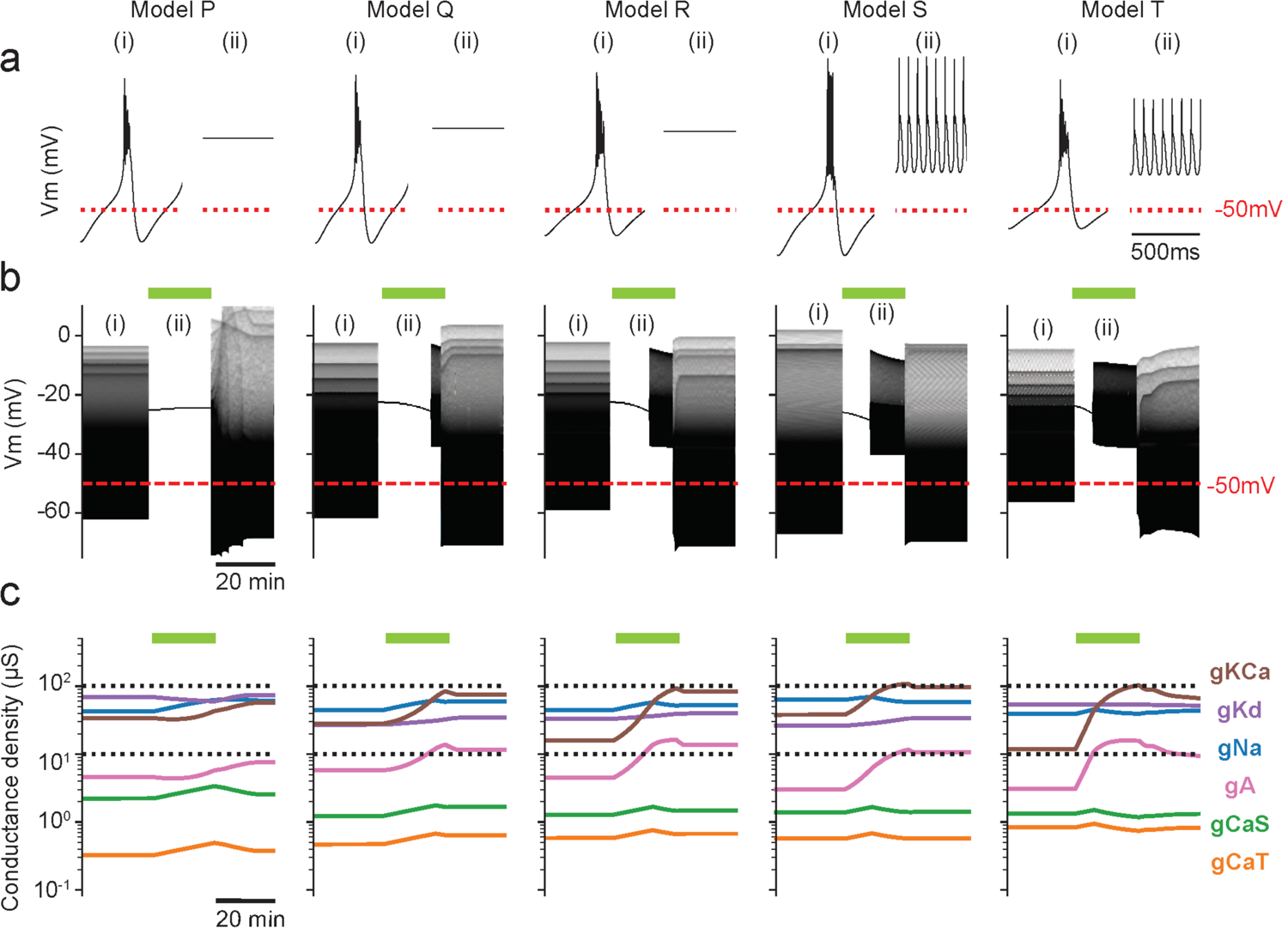
Time course of recovery depends on starting conductance densities. Response of five model bursting neurons (models P, Q, R, S and T) with different conductance densities exposed to a high potassium perturbation, represened by the green bars. **(a)** Representative traces of five models in control (*i*) and in elevated extracellular K (*ii*). **(b)** Membrane potential trace over the high potassium perturbation. All models become quiescent upon perturbation and recover spiking over a variable amount of time. **(c)** Conductance densities for each model over time.

To investigate the reasons behind this variability, we plotted the evolution of each of the conductances for the five models in Figure 5c. The recovery mechanism in each model responds differently to the same perturbation because in each case the neuron must regain spiking activity starting from a different point in conductance space. Across all models, note that the specific conductance changes in response to high potassium differ, and that in all cases the potassium conductances increase. This increase in potassium conductances makes intuitive sense, as the high potassium perturbation has the effect of reducing the driving force for all potassium currents in the model neuron.

Hence, a subsequent increase in total potassium conductance might bring the neuron closer to the baseline activity state. In Figure 5c, the H conductance is not shown because g_H_ < 10^−2^µS.

### Models with different conductance densities all retain robustness to high potassium saline, but specific changes in currents and recovery patterns vary

For each of the 9 of models described above we simulated the entire experiment of four high potassium applications, including the long wash period between applications three and four (Figs. 3, 4). Despite the variability in time to recovery in the first high potassium application (Fig. 5), all models regained spiking activity in high potassium saline and retained this enhanced robustness over subsequent applications of high potassium saline. Figure 6 shows the membrane potential of two representative models (model Q – Fig. 6a and model T, Fig. 6b) over the entire experiment. To visualize the differences in current contribution and dynamics, we plotted the currentscapes^50^ for each model below the voltage traces at some time stamps of interest: baseline (Fig. 6*i*), 10 minutes in first high potassium application (Fig. 6*ii*), 10 minutes in the fourth high potassium application (Fig. 6*iii*) and in the final wash (Fig. 6*iv*). Because the initial conductances are different for these two models, so are the contributions of each current to the baseline activity. In model Q the control activity shows a sizeable contribution of I_&_, (Fig. 6a*i*) but in model T, I_&_ is negligible and is replaced by larger contributions of I_CaT_, I_CaS_ and leak (Fig. 6b*i*) The A current, I_-_, contributes substantially to the activity in model Q (Fig. 6b*i*) but its contribution in model T (Fig. 6b*i*) is small. In response to the first high potassium perturbation, both models become quiescent but model T (Fig. 6b) recovers spiking more quickly (Fig. 6b*ii*).

**Figure 6.**
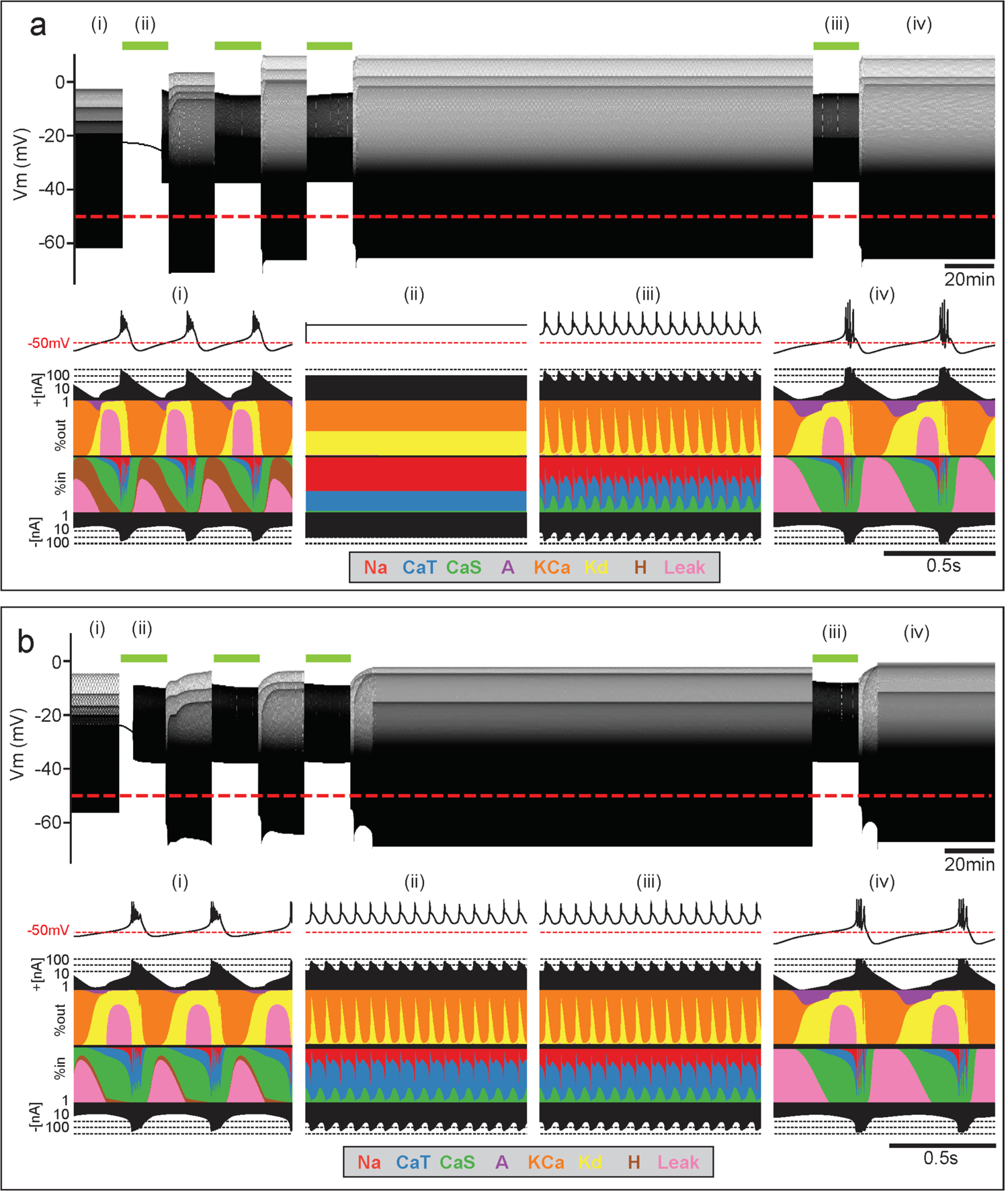
Models with different starting conductance densities retain robustness to high potassium saline, but specific changes in currents and recovery patterns vary. The green bars above the voltage traces represent the time of high potassium perturbation **(a)** The response of model Q to the entire high potassium experiment. **(b)** The response of model T to the entire high potassium experiment. Timepoints of interest are consistent between models-baseline (*i*), 10 minutes in first high potassium application (*ii*), 10 minutes in the fourth high potassium application (*iii*) and final wash (*iv*). **Top panels:** Membrane potential over time. **Bottom panels:** Currentscapes (*i-iv*). The colored panels show the percentage contribution of each individual current to the total inward or outward current over time. The black filled curves on the top and bottom indicate total inward or outward currents respectively on a logarithmic scale.

## Discussion

Neurons are long-lived cells that must perform reliably to ensure an animal’s survival. However, their components such as ion channels and synaptic proteins last days to weeks and must be constantly replaced. Thus, nervous systems are both blessed and cursed with flexibility. To maintain stable function and respond appropriately to changes in the environment, neurons and neural circuits can adapt and change over timescales ranging from milliseconds to a lifetime. Therefore, there are a plethora of activity-dependent mechanisms that regulate neuronal excitability.

Many studies focus on homeostatic mechanisms in neural circuits involving changes in gene expression and insertion of new ion channels into the membrane, typically occurring over hours to days^12, 18, 42, 51, 52^. However, there are also many examples of faster adaptation, sometimes described as rapid homeostatic plasticity^29, 38, 53^. Very rapid plasticity on the order of milliseconds to seconds, such as spike frequency adaptation or facilitation can arise from ion channel properties. These processes are critical for shaping neuronal responses, and may play a role in shaping working memory, signal transduction and many behaviors^54, 55^. Activity-dependent changes in excitability can also occur on the timescale of several minutes^20^, too long to depend on the kinetics of ion channels. On these timescales, changes in effective conductance densities can occur through calcium-dependent signaling cascades leading to phosphorylation or insertion of ion channels^26, 27, 29^.

These different activity-dependent processes, occurring on different timescales, have often been classified and segregated accordingly. However, real neurons must transition between multiple adaptation mechanisms seamlessly. This study highlights a bridge between timescales of adaptation in which rapid activity-dependent adaptation to global perturbation is retained long-term.

### Rapid and long-lasting adaptation in a conductance-based neuron model

In this study, we describe rapid adaptation in pyloric neurons following global depolarization by high potassium; this adaptation has a long-lasting effect on the circuit and affects the neuron’s response to future high potassium applications although the baseline activity appears unchanged in control saline. To better understand how this could occur, we evaluated several computational models with different properties and their response to a high potassium perturbation. Importantly, the model of a conventional conductance-based model failed to recover spiking activity in high potassium saline (Fig. 4a). This suggests that the very rapid adaptation processes determined by ion channel properties are not sufficient to account for the response of neurons to high potassium saline. Therefore, we turned to models of homeostatic plasticity which allow conductance densities to change in an activity-dependent fashion. Recent models of homeostatic plasticity link activity-dependent changes in internal calcium concentrations to changes in channel mRNA and thus conductance densities^23, 31, 33^. But changes in conductance need not rely on relatively slow changes in gene expression and subsequent translation. Here we considered how rapid changes to effective conductance density could also be activity-dependent and change the long-term excitability of a model neuron.

We propose two biologically plausible modifications to an existing homeostatic model^30^ that allow for rapid, long-lasting adaptation to perturbation while preserving normal baseline activity. This model implements three sensors that monitor the calcium current over different timescales. Specifically, each calcium sensor can be thought to represent a calcium-dependent process in the cell with different dynamics. For example, the three sensors could each represent a different calcium binding protein. In the original formulation of the model^30^, if any of the three sensors diverges from its set-point, conductance densities in the model neuron will change until the sensor returns to its calcium target. One modification we made is to assume that neuronal conductances cannot grow infinitely. Aside from obvious physical limitations on the number of ion channels that a neuron can contain, neurons may also limit their maximum conductance density to balance appropriate signaling with energy efficiency^56, 57^. This modification allows us to create model neurons with three calcium sensors whose target activity is stable over long periods of time^23^.

The most salient modification we made is to implement a gating mechanism that combines the readings of the three sensors to turn the homeostatic regulation of effective conductances on or off. Importantly, this new rule allows a model neuron to escape the requirement that all three sensors be satisfied and can turn off homeostatic regulation when target activity is “good enough”. This is in keeping with numerous studies demonstrating that many neuronal properties such as ion channel composition, synaptic weights and dendrite morphology can be sloppily tuned, and neurons can still function properly^58–60^. Biologically, the linking of the three calcium filters in the model could represent interactions between the different calcium binding processes within a cell. Given the vast complexity of calcium signaling processes, it would be unsurprising if multiple calcium binding processes were needed to initiate changes in effective channel conductance. Investigation into the specific signaling cascades or biological determinants of this adaptation are a topic for future experimental investigation.

Our results establish that using this feedback system, long-lasting adaptation to diverse perturbations and stimuli can be achieved in model neurons. Because the scheme does not require that neurons return to an exact equilibrium point, the model neurons can now retain a trace memory of past experiences even when they return to normal baseline activity. This sort of adaptation can result in degenerate circuits. For instance, models with identical starting conductances can acquire different neuronal properties despite maintaining similar activity patterns, depending on the perturbations each model is subject to. The gating scheme also opens the possibility of having more freedom in the way conductances are modified. Undoubtedly there will be rules that will be more efficient at recovering from specific perturbations, and this study provides a framework for activity-dependent models that can recover from any number of challenges, much like biological neurons. The ability to flexibly change the feedback rules would guarantee that if recovery is possible, it will happen if the neuron is kept in the perturbed condition for long enough.

### High potassium perturbations in experiments and medicine

The concentration of potassium both inside and outside cells is a critical component to proper physiological function. Despite this, few studies have focused on the acute and long-term effects of changing potassium levels. Our study highlights the possible consequences of even brief shocks of high potassium saline to a nervous system. Acute elevation of potassium concentrations is often used in experiments to rapidly excite or depolarize neurons as a proxy for excitatory inputs ^8–10^. Here, we demonstrate that adaptation to elevated high potassium saline can occur rapidly, and significantly change the excitability and intrinsic properties of neurons within minutes^38^. Therefore, studies using high potassium saline or other depolarizing stimuli should consider the possibility of rapid changes in neuronal excitability. Notably, adaptation acquired when neurons are stimulated with high potassium can be retained long after the perturbation has passed, even if baseline activity reverts and appears to be unchanged. Thus, long-term adaptation could have implications for a host of disease states involving repeated insults associated with high extracellular potassium. This phenomenon could be particularly important for understanding the long-term effects of epileptic seizures and kindling of localized seizures^61^. Within a seizure locus, extracellular potassium levels rapidly increase^62, 63^. Neurons experiencing this perturbation may change their conductance densities in response, and these changes may be maintained even after activity returns to normal levels. This sort of adaptation could exacerbate or ameliorate the severity of repeated seizures in the same locus. Similarly, these dynamics have been shown to affect peripheral nerves in patients with chronic kidney disease^7^.

### Persistent, cryptic memory in neurons following perturbation

Theoretical and experimental evidence shows that seemingly identical activity patterns in neurons can arise from widely variable underlying parameters ^33, 44, 49, 50, 53, 59, 64–67^. It has been observed that individual differences between human patients lead to different outcomes in cases of stroke^68, 69^ and traumatic brain injury^70, 71^. Similarly, the pyloric rhythm of the STG responds stereotypically and robustly to many perturbations including temperature^35, 72^ and pH^36, 37^ within a permissive range; outside this universal permissive range, each individual circuit can be more or less robust to a given perturbation and is disrupted in a unique way^42, 44, 79^. The pyloric rhythm is also variable in its response to high potassium saline^38^ (Fig. 1C), and this variability likely arises from different initial conductance densities (Fig. 5). In all these cases, individual variability between circuits is invisible at baseline conditions and only revealed by a critical perturbation.

The origin of individual variability in neuronal circuits is a topic of ongoing exploration and debate ^53, 66, 73^. Here we show that neurons can rapidly adapt to changes in the environment without maintaining precise levels of any given conductance. An interesting suggestion of this study is that circuits may evolve over time in response to environmental perturbations, while retaining their normal physiological function. Here, we show rapid adaptation to a high potassium perturbation in both biological and model neurons where the activity pattern returns to the baseline state after the perturbation is removed. We show that even though baseline neuronal activity appears unchanged, the robustness of neurons to future perturbation is altered. In this way, past exposure to high potassium saline acts as a prior, e.g. a past experience will bias the outcome of a future output^74–76^. Hence, adaptation in response to perturbation can be long-lasting and invisible when observing only baseline activity.

## Acknowledgments

We thank Dr. Jonathan Touboul for useful discussions, and Janis Li for assistance running biological experiments. This work was supported by NIH R35 NS097343 (E.M., L.A.) and F31-NS113383 (M.R.).

## Author Contributions

M.R. and L.A. contributed equally to this work. M.R., L.A. and E.M. conceived the study design. M.R. performed biological experiments. L.A. performed the modeling experiments. All authors discussed the results and contributed to writing and editing the final manuscript and figures.

## Data availability

The data reported in this manuscript are available from the corresponding author upon reasonable request.

## Code availability

The MATLAB analysis code and Python simulation code reported in this manuscript is available at the Marder lab GitHub (https://github.com/marderlab) upon publication.

## Methods

### Animals and dissections

Adult male Jonah Crabs, *Cancer borealis*, (N = 20) were obtained from Commercial Lobster (Boston, MA) from January to August 2020 and maintained in artificial seawater at 10-12°C in a 12-hour light/dark cycle. On average, animals were acclimated in the laboratory for one week before use. Prior to dissection, animals were placed on ice for at least 30 minutes. Dissections were performed as previously described^1^. The stomach was dissected from the animal and the intact stomatogastric nervous system (STNS) was removed from the stomach including the commissural ganglia, esophageal ganglion and stomatogastric ganglion (STG) with connecting motor nerves. The STNS was pinned in a Sylgard-coated (Dow Corning) dish and continuously superfused with 11°C saline.

### Solutions

Physiological (control) *Cancer borealis* saline was composed of 440 mM NaCl, 11 mM KCl, 26 mM MgCl_2_, 13 mM CaCl_2_, 11 mM Trizma base, 5 mM maleic acid, pH 7.4-7.5 at 23°C (approximately 7.7-7.8 pH at 11°C). High [K^+^] saline (2.5x[K^+^], 27.5mM KCl) was prepared by adding more KCl salt to the normal saline.

### Electrophysiology

Intracellular recordings from STG somata were made in the desheathed STG with 10–30 MΩ sharp glass microelectrodes filled with internal solution: 10 mM MgCl_2_, 400 mM potassium gluconate, 10 mM HEPES buffer, 15 mM NaSO_4_, 20 mM NaCl^2^. Intracellular signals were amplified with an Axoclamp 900A amplifier (Molecular Devices, San Jose).

Extracellular nerve recordings were made by building wells around nerves using a mixture of Vaseline and mineral oil and placing stainless-steel pin electrodes within the wells to monitor spiking activity. Extracellular nerve recordings were amplified using model 3500 extracellular amplifiers (A-M Systems). Data were acquired using a Digidata 1440 digitizer (Molecular Devices, San Jose) and pClamp data acquisition software (Molecular Devices, San Jose, version 10.5). For identification of Pyloric Dilator (PD) neurons, somatic intracellular recordings were matched to extracellular action potentials on the pyloric dilator nerve (*pdn*) and/or the lateral ventricular nerve (*lvn*).

### Elevated [K^+^] saline application

For all preparations, baseline activity of the PD neuron was first recorded for 30 minutes in control saline. Following the baseline recording, the STNS was superfused with 2.5x[K^+^] saline for 20 minutes, followed by a 20-minute wash in control saline. This pattern was repeated, alternating between 20 minute 2.5x[K^+^] saline and physiological control saline three times. In some experiments, the preparation was then washed in physiological saline for three hours before a final fourth 20-minute 2.5x[K^+^] saline application and a final 20-minute wash.

### Data acquisition and analysis

Recordings were acquired using Clampex software (pClamp Suite by Molecular Devices, San Jose, version 10.5) and visualized and analyzed using custom MATLAB analysis scripts. These scripts were used to detect and measure voltage response amplitudes and membrane potentials, plot raw recordings and processed data, generate raster plots, and perform some statistical analyses.

### Analysis of interspike interval distributions

To extract spike times, we used a custom spike identification and sorting software (called “crabsort”) which uses a TensorFlow based machine-learning algorithm.

Crabsort is freely available at https://github.com/sg-s/crabsort and its use is described in Powell et al. (2021)^3^. Distributions of inter-spike intervals (ISIs) were calculated within 2-minute bins. Hartigan’s dip test of unimodality^4^ was used to obtain the dip statistic for each of these distributions. This dip statistic was compared to Table 1 in Hartigan and Hartigan^4^ to find the probability of multi-modality. The test creates a unimodal distribution function that has the smallest value deviations from the experimental distribution function. The largest of these deviations is the dip statistic. The dip statistic shows the probability of the experimental distribution function being bimodal. Larger value dips indicate that the empirical data are more likely to have multiple modes^4^. For visualizing spiking activity in raster plots, if the dip statistic was 0.05 or higher the neuron was considered to be bursting. If the dip statistic was lower than 0.05 the neuron was considered to be tonically firing. In neurons with less than 30 action potentials per minute, there were too few spikes to calculate an accurate dip statistic and the neurons are labeled as tonically firing.

**TABLE I.** Control scheme used in (Liu et al., 1998)

|  | Na | CaS | CaT | Kd | KCa | A | H |
| --- | --- | --- | --- | --- | --- | --- | --- |
| A | 1 | 0 | 0 | 1 | 0 | 0 | 0 |
| B | 0 | 1 | 1 | -1 | -1 | -1 | 1 |
| C | 0 | 0 | 0 | 0 | -1 | -1 | 1 |

### Computational modeling of bursting neurons

In this work we implemented modifications of the model by Liu et al. (1998). The model neuron has a sodium current, I_Na_; transient and slow calcium currents I_CaT_ and I_CaS_; a transient potassium current, I_A_; a calcium dependent potassium current, I_KCa_; a delayed rectifier potassium current, I_Kd_; a hyperpolarization-activated inward current, I_H_; and a leak current I_Leak_. The model uses its calcium currents to modify its conductance densities to achieve a target activity. The model has three sensors that monitor the calcium currents over different time scales and are named accordingly as fast (F), slow (S) and dc (D). The activity of these sensors are used to drive changes in the maximal conductances using the following equation,

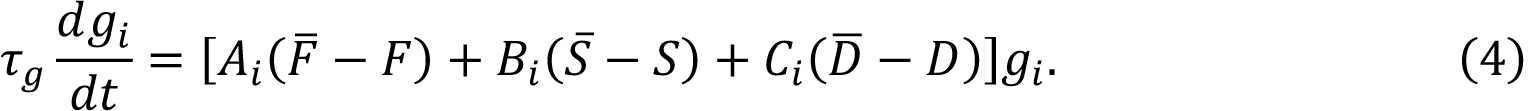

Here F^–^, S^–^ and C^–^ are *target values* for the average activity of the sensors and τ_g_ is the time scale of conductance evolution and index *i* specifies the current type. The coefficients *A_i_*, *B_i_* and *C_i_* determine what the model will do with each conductance when the average activity of the corresponding sensor is off-target. Hereafter, we refer to these coefficients as the “control scheme” or “scheme”. The scheme used by Liu et al. (1998) is reproduced in table IIIA.

We can rewrite the equations in vector notation by introducing the maximal conductance vector **g** = {g_i_} with g_i_. the maximal conductance of channel type *i* and error vector δ as follows,

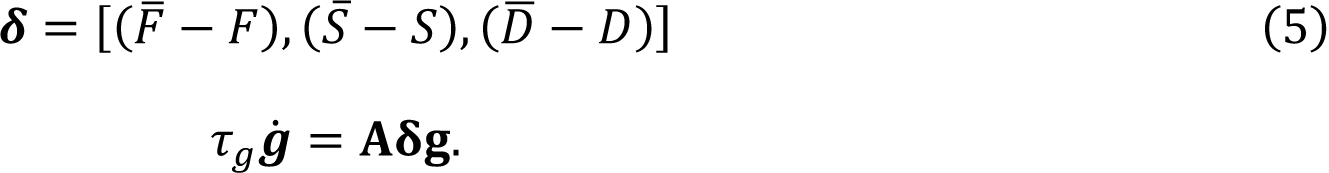

In this notation the control scheme in table IIIA is represented by a matrix **A** and the distance between each sensor and its target is represented by vector δ(t).

We added a cubic term in each component of ġ to prevent the model’s conductances from growing exponentially large. We found that there is a range of values of γ for which the model neuron always settles into a periodic bursting regime. For Figure 4b we used γ_i_ = 10^5^ for all currents except I_A_ where we used γ_0_ = 60 × 10^−5^

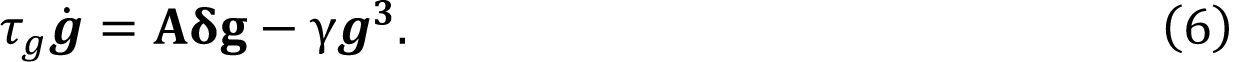

To modulate the timescale of conductance change, τ_-_, we defined a feedback signal *S*_f_ as follows,

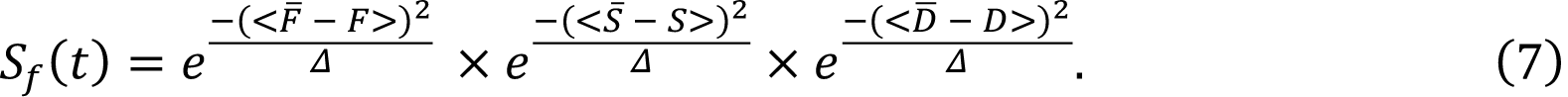

This quantity is the product of three gaussian functions that will take values close to 1 if the corresponding sensor is near its set point and values close to 0 otherwise. By definition, S_3_ takes values close to 1 if all three sensors are near their targets at the same time, and close to 0 otherwise. Parameter Δ determines how close the sensors need to be to their set points to produce a high feedback. In this work we set Δ = 0.001.

The timescale for evolution of conductance densities is modulated by a state variable α as follows,

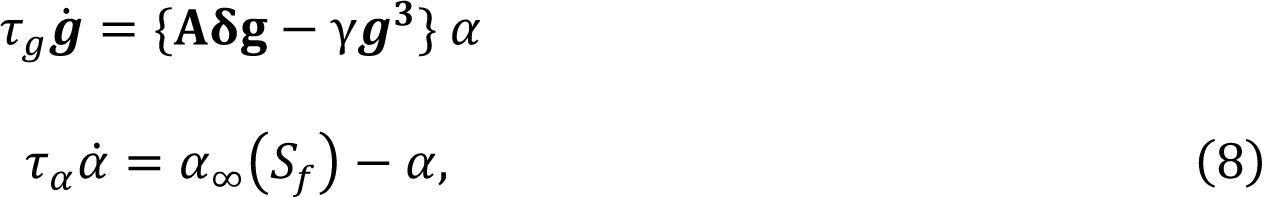

with

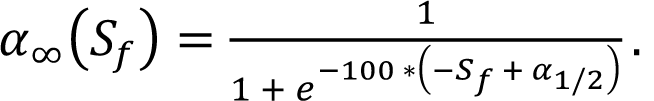

The parameters in function α_A_ were chosen so that α_A_ ≈ 0 if S_3_ > 0.2 and α_A_(S_3_) ≈ 1 otherwise. In this way high feedback switches α(t) → 0 over a timescale τ_@_ = 1000 *msec*. In this equation, α_B/F_ is the half-maximal activation of α. We set α_B/F_ = 0.075 to allow gating of conductance regulation. Notice that using this parameter, we can switch between models with and without conductance regulation. If α_B/F_ = −1, α → 0 and there is no gating. If α_B/F_ = 10, then α → 1 and the regulation mechanism is always on.

We simulated the application of high potassium saline in the models by changing the equilibrium potential, E_K_, of the potassium currents from −80mV (control) to −40mV (high potassium). In addition, we changed the reversal potential of the leak conductance from −50mV (control) to −40mV (high potassium) because the leak current is a non-specific cation current with a sizable potassium contribution.

All the equations and parameters of the sensors, and the activation functions are identical to those in Liu et al. (1998). The models were simulated using an exponential-Euler scheme with step dt = 0.1 msec ^5^. All simulations were performed in commercially available computers using python. Code to reproduce the simulations is available upon request.

### Generating multiple models

We obtained multiple models by simulating equation (3) starting from small random initial conductances and allowing them to evolve under control conditions until they settled into their target bursting regimes.

## Statistics

Statistical analysis and plotting were carried out using MATLAB 2020b built in functions for all analyses as described above. All electrophysiology analysis scripts are available at the Marder lab GitHub (https://github.com/marderlab).

## Work Cited

1. Baylor, D.A. & Nicholls, J.G. Changes in extracellular potassium concentration produced by neuronal activity in the central nervous system of the leech. J. Physiol. 203, 555–569 (1969).

2. Chauvette, S., Soltani, S., Seigneur, J. & Timofeev, I. *In vivo* models of cortical acquired epilepsy. J. Neurosci. Methods 260, 185–201 (2016).

3. Katayama, Y., Becker, D.P., Tamura, T. & Hovda, D.A. Massive increases in extracellular potassium and the indiscriminate release of glutamate following concussive brain injury. J. Neurosurg. 73, 889–900 (1990).

4. Rodgers, C.I., et al. Stress preconditioning of spreading depression in the locust CNS. PLoS One 2, e1366 (2007).

5. Morrison III, B., Elkin, B.S., Dollé, J.-P. & Yarmush, M.L. *In vitro* models of traumatic brain injury. Annu. Rev. Biomed. Eng. 13, 91–126 (2011).

6. Seeburg, D.P. & Sheng, M. Activity-induced Polo-like kinase 2 is required for homeostatic plasticity of hippocampal neurons during epileptiform activity. J. Neurosci. 28, 6583–6591 (2008).

7. Arnold, R., et al. Evidence for a causal relationship between hyperkalaemia and axonal dysfunction in end-stage kidney disease. Clin. Neurophysiol. 125, 179–185 (2014).

8. Sharma, N., Gabel, Harrison W. & Greenberg, Michael E. A Shortcut to Activity-Dependent Transcription. Cell 161, 1496–1498 (2015).

9. Rybak, I.A., Molkov, Y.I., Jasinski, P.E., Shevtsova, N.A. & Smith, J.C. Rhythmic Bursting in the Pre-Bötzinger Complex. in *Prog*. Brain Res. 1–23 (Elsevier, 2014).

10. Ballerini, L., Galante, M., Grandolfo, M. & Nistri, A. Generation of rhythmic patterns of activity by ventral interneurones in rat organotypic spinal slice culture. J. Physiol. 517, 459–475 (1999).

11. Ruangkittisakul, A., Panaitescu, B. & Ballanyi, K. K^+^ and Ca^2+^ dependence of inspiratory-related rhythm in novel “calibrated” mouse brainstem slices. Respir. Physiol. Neurobiol. 175, 37–48 (2011).

12. O’Leary, T., van Rossum, M.C. & Wyllie, D.J. Homeostasis of intrinsic excitability in hippocampal neurones: dynamics and mechanism of the response to chronic depolarization. J. Physiol. 588, 157–170 (2010).

13. Grubb, M.S. & Burrone, J. Activity-dependent relocation of the axon initial segment fine-tunes neuronal excitability. Nature 465, 1070–1074 (2010).

14. Rannals, M.D. & Kapur, J. Homeostatic strengthening of inhibitory synapses is mediated by the accumulation of GABAA receptors. J. Neurosci. 31, 17701–17712 (2011).

15. Turrigiano, G. Homeostatic synaptic plasticity: local and global mechanisms for stabilizing neuronal function. Cold Spring Harb. Perspect. Biol. 4, a005736 (2012).

16. Turrigiano, G.G., Leslie, K.R., Desai, N.S., Rutherford, L.C. & Nelson, S.B. Activity-dependent scaling of quantal amplitude in neocortical neurons. Nature 391, 892–896 (1998).

17. Turrigiano, G.G. & Nelson, S.B. Homeostatic plasticity in the developing nervous system. Nat. Rev. Neurosci. 5, 97–107 (2004).

18. Temporal, S., Lett, K.M. & Schulz, D.J. Activity-dependent feedback regulates correlated ion channel mRNA levels in single identified motor neurons. Curr. Biol. 24, 1899–1904 (2014).

19. Brickley, S.G., Revilla, V., Cull-Candy, S.G., Wisden, W. & Farrant, M. Adaptive regulation of neuronal excitability by a voltage-independent potassium conductance. Nature 409, 88–92 (2001).

20. Golowasch, J., Abbott, L.F. & Marder, E. Activity-dependent regulation of potassium currents in an identified neuron of the stomatogastric ganglion of the crab *Cancer borealis*. J. Neurosci. 19, RC33-RC33 (1999).

21. Desai, N.S. Homeostatic plasticity in the CNS: synaptic and intrinsic forms. J. Physiol. (Paris) 97, 391–402 (2003).

22. Marder, E. & Goaillard, J.-M. Variability, compensation and homeostasis in neuron and network function. Nat. Rev. Neurosci. 7, 563–574 (2006).

23. O’Leary, T., Williams, A.H., Franci, A. & Marder, E. Cell types, network homeostasis, and pathological compensation from a biologically plausible ion channel expression model. Neuron 82, 809–821 (2014).

24. Mease, R.A., Famulare, M., Gjorgjieva, J., Moody, W.J. & Fairhall, A.L. Emergence of adaptive computation by single neurons in the developing cortex. J. Neurosci. 33, 12154–12170 (2013).

25. Hengen, K.B., Lambo, M.E., Van Hooser, S.D., Katz, D.B. & Turrigiano, G.G. Firing rate homeostasis in visual cortex of freely behaving rodents. Neuron 80, 335–342 (2013).

26. Misonou, H., et al. Regulation of ion channel localization and phosphorylation by neuronal activity. Nat. Neurosci. 7, 711–718 (2004).

27. Park, K.-S., Mohapatra, D.P., Misonou, H. & Trimmer, J.S. Graded regulation of the Kv2. 1 potassium channel by variable phosphorylation. Science 313, 976-979 (2006).

28. Capera, J., Serrano-Novillo, C., Navarro-Pérez, M., Cassinelli, S. & Felipe, A. The potassium channel odyssey: mechanisms of traffic and membrane arrangement. Int. J. Mol. Sci. 20, 734 (2019).

29. Frank, C.A., Kennedy, M.J., Goold, C.P., Marek, K.W. & Davis, G.W. Mechanisms underlying the rapid induction and sustained expression of synaptic homeostasis. Neuron 52, 663–677 (2006).

30. Liu, Z., Golowasch, J., Marder, E. & Abbott, L. A model neuron with activity-dependent conductances regulated by multiple calcium sensors. J. Neurosci. 18, 2309–2320 (1998).

31. Gorur-Shandilya, S., Marder, E. & O’Leary, T. Activity-dependent compensation of cell size is vulnerable to targeted deletion of ion channels. Scientific Reports 10 (2020).

32. Golowasch, J., Goldman, M.S., Abbott, L.F. & Marder, E. Failure of averaging in the construction of a conductance-based neuron model. J. Neurophysiol. 87, 1129–1131 (2002).

33. O’Leary, T. & Marder, E. Temperature-Robust Neural Function from Activity-Dependent Ion Channel Regulation. Curr. Biol. 26, 2935–2941 (2016).

34. LeMasson, G., Marder, E. & Abbott, L. Activity-dependent regulation of conductances in model neurons. Science 259, 1915–1917 (1993).

35. Haddad, S.A. & Marder, E. Circuit robustness to temperature perturbation is altered by neuromodulators. Neuron 100, 609–623 (2018).

36. Haley, J.A., Hampton, D. & Marder, E. Two central pattern generators from the crab, *Cancer borealis*, respond robustly and differentially to extreme extracellular pH. eLife 7, e41877 (2018).

37. Ratliff, J., Franci, A., Marder, E. & O’Leary, T. Neuronal oscillator robustness to multiple global perturbations. Biophys. J. (2021).

38. He, L.S., et al. Rapid adaptation to elevated extracellular potassium in the pyloric circuit of the crab, Cancer borealis. J. Neurophysiol. 123, 2075–2089 (2020).

39. Harris-Warrick, R.M., Marder, E., Selverston, A.I. & Moulins, M. Dynamic biological networks: the stomatogastric nervous system (MIT press, 1992).

40. Soofi, W., et al. Phase maintenance in a rhythmic motor pattern during temperature changes *in vivo*. J. Neurophysiol. 111, 2603–2613 (2014).

41. Morris, L.G. & Hooper, S.L. Muscle response to changing neuronal input in the lobster (Panulirus interruptus) stomatogastric system: spike number-versus spike frequency-dependent domains. J. Neurosci. 17, 5956–5971 (1997).

42. Turrigiano, G., Abbott, L.F. & Eve, M. Activity-Dependent Changes in the Intrinsic Properties of Cultured Neurons. *Science*, New Series 264, 974–977 (1994).

43. O’Leary, T. & Marder, E. Mapping Neural Activation onto Behavior in an Entire Animal. Science 344, 372–373 (2014).

44. Schulz, D.J., Goaillard, J.-M. & Marder, E. Variable channel expression in identified single and electrically coupled neurons in different animals. Nat. Neurosci. 9, 356–362 (2006).

45. Goaillard, J.-M., Taylor, A.L., Schulz, D.J. & Marder, E. Functional consequences of animal-to-animal variation in circuit parameters. Nat. Neurosci. 12, 1424–1430 (2009).

46. Ransdell, J.L., Faust, T.B. & Schulz, D.J. Correlated Levels of mRNA and Soma Size in Single Identified Neurons: Evidence for Compartment-specific Regulation of Gene Expression. Front. Mol. Neurosci. 3, 116 (2010).

47. Schulz, D.J., Goaillard, J.M. & Marder, E.E. Quantitative expression profiling of identified neurons reveals cell-specific constraints on highly variable levels of gene expression. Proc. Natl. Acad. Sci. U.S.A. 104, 13187–13191 (2007).

48. Tobin, A.E., Cruz-Bermudez, N.D., Marder, E. & Schulz, D.J. Correlations in ion channel mRNA in rhythmically active neurons. PLoS One 4, e6742 (2009).

49. Prinz, A.A., Bucher, D. & Marder, E. Similar network activity from disparate circuit parameters. Nat. Neurosci. 7, 1345–1352 (2004).

50. Alonso, L.M. & Marder, E. Visualization of currents in neural models with similar behavior and different conductance densities. eLife 8, e42722 (2019).

51. Turrigiano, G. Homeostatic synaptic plasticity: local and global mechanisms for stabilizing neuronal function. Cold Spring Harb. Perspect. Biol. 4, a005736 (2012).

52. Desai, N.S., Rutherford, L.C. & Turrigiano, G.G. Plasticity in the intrinsic excitability of cortical pyramidal neurons. Nat. Neurosci. 2, 515–520 (1999).

53. Lane, B.J., Samarth, P., Ransdell, J.L., Nair, S.S. & Schulz, D.J. Synergistic plasticity of intrinsic conductance and electrical coupling restores synchrony in an intact motor network. eLife 5, e16879 (2016).

54. Roach, J.P., Sander, L.M. & Zochowski, M.R. Memory recall and spike-frequency adaptation. Phys. Rev. 93, 052307 (2016).

55. Benda, J., Longtin, A. & Maler, L. Spike-frequency adaptation separates transient communication signals from background oscillations. J. Neurosci. 25, 2312–2321 (2005).

56. Sengupta, B., Faisal, A.A., Laughlin, S.B. & Niven, J.E. The effect of cell size and channel density on neuronal information encoding and energy efficiency. J. Cereb. Blood Flow Metab. 33, 1465–1473 (2013).

57. Kim, M., McKinnon, D., MacCarthy, T., Rosati, B. & McKinnon, D. Regulatory evolution and voltage-gated ion channel expression in squid axon: selection-mutation balance and fitness cliffs. PLoS One 10, e0120785 (2015).

58. Otopalik, A.G., Sutton, A.C., Banghart, M. & Marder, E. When complex neuronal structures may not matter. eLife 6 (2017).

59. Sakurai, A., Tamvacakis, A.N. & Katz, P.S. Hidden synaptic differences in a neural circuit underlie differential behavioral susceptibility to a neural injury. eLife 3, e02598 (2014).

60. Marder, E., Goeritz, M.L. & Otopalik, A.G. Robust circuit rhythms in small circuits arise from variable circuit components and mechanisms. Curr. Opin. Neurobiol. 31, 156–163 (2015).

61. Chauvette, S., Soltani, S., Seigneur, J. & Timofeev, I. In vivo models of cortical acquired epilepsy. J. Neurosci. Methods 260, 185–201 (2016).

62. Moody, W.J., Jr., Futamachi, K.J. & Prince, D.A. Extracellular potassium activity during epileptogenesis. Exp. Neurol. 42, 248–263 (1974).

63. Fröhlich, F., Bazhenov, M., Iragui-Madoz, V. & Sejnowski, T.J. Potassium dynamics in the epileptic cortex: new insights on an old topic. Neuroscientist 14, 422–433 (2008).

64. Klassen, T., et al. Exome sequencing of ion channel genes reveals complex profiles confounding personal risk assessment in epilepsy. Cell 145, 1036–1048 (2011).

65. Fuzik, J., et al. Integration of electrophysiological recordings with single-cell RNA-seq data identifies neuronal subtypes. Nat. Biotechnol. 34, 175 (2016).

66. Ciarleglio, C.M., et al. Correction: Multivariate analysis of electrophysiological diversity of Xenopus visual neurons during development and plasticity. eLife 5, e14282 (2016).

67. Goldman, M.S., Golowasch, J., Marder, E. & Abbott, L. Global structure, robustness, and modulation of neuronal models. J. Neurosci. 21, 5229–5238 (2001).

68. Price, C.J., Seghier, M.L. & Leff, A.P. Predicting language outcome and recovery after stroke: the PLORAS system. Nat. Rev. Neurol. 6, 202–210 (2010).

69. Cramer, S.C. Repairing the human brain after stroke: I. Mechanisms of spontaneous recovery. Ann. Neurol. 63, 272–287 (2008).

70. Juengst, S.B., Terhorst, L., Kew, C.L. & Wagner, A.K. Variability in daily self-reported emotional symptoms and fatigue measured over eight weeks in community dwelling individuals with traumatic brain injury. Brain injury 33, 567–573 (2019).

71. Dams-O’Connor, K., et al. Rehospitalization over 10 years among survivors of TBI: A National Institute on Disability, Independent Living and Rehabilitation Research (NIDILRR) Traumatic Brain Injury Model Systems Study. J. Head Trauma Rehabil. 32, 147 (2017).

72. Tang, L.S., Taylor, A.L., Rinberg, A. & Marder, E. Robustness of a rhythmic circuit to short-and long-term temperature changes. J. Neurosci. 32, 10075–10085 (2012).

73. Nelson, S.B. & Turrigiano, G.G. Strength through diversity. Neuron 60, 477–482 (2008).

74. Verstynen, T. & Sabes, P.N. How each movement changes the next: an experimental and theoretical study of fast adaptive priors in reaching. J. Neurosci. 31, 10050–10059 (2011).

75. Yang, J., Lee, J. & Lisberger, S.G. The interaction of Bayesian priors and sensory data and its neural circuit implementation in visually guided movement. J. Neurosci. 32, 17632–17645 (2012).

76. Darlington, T.R., Tokiyama, S. & Lisberger, S.G. Control of the strength of visual-motor transmission as the mechanism of rapid adaptation of priors for Bayesian inference in smooth pursuit eye movements. J. Neurophysiol. 118, 1173–1189 (2017).

## Work Cited

77. Gutierrez, G.J. & Grashow, R.G. Cancer borealis stomatogastric nervous system dissection. J. Vis. Exp. (JoVE) (2009).

78. Hooper, S.L., Thuma, J.B., Guschlbauer, C., Schmidt, J. & Büschges, A. Cell dialysis by sharp electrodes can cause nonphysiological changes in neuron properties. J. Neurophysiol. 114, 1255–1271 (2015).

79. Powell, D., Haddad, S.A., Gorur-Shandilya, S. & Marder, E. Coupling between fast and slow oscillator circuits in Cancer borealis is temperature-compensated. eLife 10, e60454 (2021).

80. Hartigan, J.A. & Hartigan, P.M. The dip test of unimodality. Ann. Stat. 13, 70–84 (1985).

81. Dayan, P. & Abbott, L.F. Theoretical neuroscience: computational and mathematical modeling of neural systems. J. Cognit. Neurosci. 15, 154–155 (2003).

